# Comparative genomics reveals a genomic and functional continuum among secondary replicons in Halobacteriota

**DOI:** 10.64898/2026.09.22.753402

**Authors:** Angélica Jara-Servin, Macarena Toll-Riera

**Affiliations:** Institut de Biologia Evolutiva (CSIC-UPF), Barcelona, Spain

## Abstract

A significant fraction of archaeal genomes have their genomic content distributed across two or more DNA molecules, yet the characteristics and evolution of these different replicons remain poorly understood. Here, we investigated the genomic and functional organization of secondary replicons in Halobacteriota using 211 complete genomes, comprising 766 replicons, including 211 primary chromosomes (chr1), and 555 secondary replicons. Comparative analyses of genomic features, including replicon size, GC content, and codon usage, revealed a continuous distribution of secondary replicons from plasmid-like to chromosome-like characteristics. Larger secondary replicons showed progressively more chromosome-like genomic and functional properties, including greater protein-family overlap with chr1 and a higher proportion of conserved gene families. At the phylum level, functional profiles substantially overlapped among replicon types, providing further support for a continuum rather than discrete functional classes. Larger replicons also showed increased protein-family duplication, while pangenome analysis revealed a greater proportion of conserved gene families, with accessory families most abundant at intermediate sizes and rare and singleton families declining with increasing replicon size. Together, these results support a model in which secondary replicons in Halobacteriota span a genomic and functional continuum from plasmid-like to chromosome-like states, with increasing size associated with greater conservation, gene duplication and progressively more chromosome-like gene content and functional profile.

## INTRODUCTION

Prokaryotic organisms, particularly bacteria and, to a lesser extent, archaea, are commonly conceived as unicellular microorganisms with a single chromosome and sometimes one or more plasmids. This view emerged alongside the first studies in microbial genomics in the 1960s (Cairns, 1963). However, over the years, bacteria harboring more than one chromosome started to be described (Suwanto and Kaplan, 1989), and as the number of these “exceptions” increased, it became clear that some bacterial species possess a genomic architecture comprising more than one chromosome, hence confirming the existence of multipartite bacterial genomes. Multipartite genomes consist of a primary chromosome and one or more additional large secondary replicons known as chromids or secondary chromosomes (diCenzo and Finan, 2017). It is important to emphasize that chromids or secondary chromosomes are not plasmids. While plasmids and megaplasmids are considerably smaller than the primary chromosome, typically differ in GC content, possess plasmid-like replication origins, and mainly carry accessory genes, secondary chromosomes are generally only slightly smaller than the primary chromosome and have a similar GC content, but retain plasmid-like replication origins and encode essential genes (diCenzo and Finan, 2017). It is estimated that about 10% of bacterial genomes are multipartite (Harrison et al., 2010), including members of environmentally widespread genera such as *Pseudoalteromonas*, *Vibrio* and *Burkholderia* (diCenzo and Finan, 2017; Riccardi et al., 2023). Although secondary chromosomes are common, their evolutionary origin remains unclear, and several hypotheses have been proposed to explain their emergence. One prominent hypothesis suggests that secondary chromosomes evolved from ancestral plasmids that became established within a host and progressively acquired essential genes, ultimately developing chromosome-like characteristics (Slater et al., 2009).

In Archaea, plasmids are relatively common; however, secondary chromosomes have been less described. Large plasmids are less common than smaller ones and have been identified primarily in members of the Halobacteriota (Wang et al., 2015). Secondary chromosomes have also been described in archaeal genomes, with their classification generally supported by experimental evidence demonstrating chromosome-like properties, including autonomous replication, the presence of essential genes, and stable inheritance (Norais et al., 2007; Ausiannikava et al., 2018). Notably, the distinction between large plasmids (or megaplasmids) and secondary chromosomes may be less clear in Archaea than in Bacteria, as many archaeal megaplasmids encode genes essential for host survival. For example, *Haloarcula marismortui* possesses a multipartite genome in which essential metabolic and informational functions are distributed across the primary chromosome, secondary chromosomes, and smaller replicons (Baliga et al., 2004). The evolutionary origin of megaplasmids and secondary chromosomes remains poorly understood in Archaea. In agreement with one hypothesis proposed for bacteria, it has been suggested that genetic exchange between the primary chromosome and plasmids gave rise to megaplasmids, which subsequently evolved into secondary chromosomes through prolonged coexistence and genetic exchange with the primary chromosome (Schwartz, 2009). However, there is also evidence suggesting that megaplasmids originated from an ancient inter-kingdom horizontal transfer event (Forterre et al., 2014). Given the absence of an operational definition that clearly distinguishes plasmids, megaplasmids and secondary chromosomes in archaeal genomes, a practical definition of an archaeal multipartite genome is one in which the genetic content is distributed across a primary chromosome and one or more secondary replicon, which could be either secondary chromosomes, megaplasmids or plasmids (irrespective of their size).

Genomes, whether bacterial or archaeal, exhibit genomic features that vary among species. Leveraging these differences provides an opportunity to analyze replicons and find shared traits at higher taxonomic ranks while also distinguishing replicon types within individual genomes. Several genomic features have proven informative for replicon classification, including codon usage, dinucleotide relative abundance, conjugative properties, and gene content (diCenzo and Finan, 2017). These comparative genomic signatures complement classical criteria used to distinguish the replicons within a particular genome, including replicon size, GC content, replication and partitioning systems, and differences in gene content among the primary chromosomes, secondary chromosomes, and plasmids. By combining these approaches, replicon identity can be evaluated not only based on intrinsic properties, but also within the broader genomic and evolutionary context of related organisms. Such an integrated approach provides insights into the degree of genomic and functional integration of secondary chromosomes, megaplasmids and plasmids with the host genome, their potential for independent maintenance, and the evolutionary processes underlying their emergence, diversification, and persistence.

Despite the increasing number of reported archaeal multipartite genomes, the evolutionary relationships among primary chromosomes, secondary chromosomes, and plasmids remain poorly understood. In particular, it is unclear whether large secondary replicons represent intermediate evolutionary forms between plasmids and chromosomes, and to what extent they have acquired chromsome-like properties. Moreover, the genomic features that best distinguish these replicon types remain poorly defined. Here, we address these gaps by analyzing complete multipartite genomes from the Halobacteriota, an archaeal lineage in which secondary replicons are highly frequent. This group therefore provides an ideal system for comparing replicon types both within individual genomes and across related taxa. Using comparative genomics, we integrate compositional, structural, and functional features to characterize replicon identity and the relationships among replicon types. Specifically, we investigate whether different replicon types are associated with distinct genomic signatures or instead span a continuum of plasmid- to chromosome-like characteristics, providing insights into the diversification and evolutionary integration of secondary replicons.

## MATERIALS AND METHODS

### Retrieval and processing of Halobacteriota multipartite genomes

We downloaded from RefSeq all 1,970 archaeal genomes assembled to the Complete Genome level, together with their corresponding JSONL metadata files, using the NCBI datasets package on May 4, 2026. Using the metadata contained in the JSONL files, we identified archaeal genomes harboring secondary chromosomes, plasmids or both, in spite of their length. After removing biological duplicates (i.e. genomes originated from the same biosample) and removing all genomes from other phyla, we retained a total of 211 complete multipartite genomes from the phylum Halobacteriota —according to the GTDB taxonomy—for subsequent analyses. Unless otherwise specified, all downstream analyses were performed using bash, R v4.5.1 (R Core Team, 2025), and Python v3.7.8. All detailed bioinformatic analysis, protocols, and datasets are available on GitHub (https://github.com/ajaraservin/archaea_secondary_replicons).

We reannotated all retained genomes using Prokka v1.14.5 (Seemann, 2014) to generate standardized genome annotations and locus-level information across all genomes. Among the 211 complete genomes, we identified 766 replicons. We used the replicon descriptions contained in contig headers to classify replicons into three categories: primary chromosomes, secondary chromosomes, and plasmids. Hereafter, the primary chromosome is referred to as chr1, whereas secondary chromosomes and plasmids are collectively referred to as secondary replicons, and individually as chr2 and plasmids, respectively. For downstream comparative analyses, we generated CDS and protein sequences independently using Prodigal v2.6.3 (Hyatt et al., 2010). To integrate the two annotation workflows, we constructed a correspondence table linking Prodigal protein identifiers with Prokka locus tags, genome accessions, contigs, and genomic coordinates. This correspondence table allowed protein-level analyses to be linked consistently to the corresponding genomic features throughout the analysis pipeline. This mapping enabled functional annotations generated using eggNOG-mapper and FANTASIA (see below) to be unambiguously associated with the corresponding proteins, genomes, replicons, and protein clusters. To conduct the taxonomic assignments, we used the Genome Taxonomy Database (GTDB) Release 232 (Chaumeil et al., 2020).

### Calculation and analysis of replicon features

We calculated length and GC content for each replicon with seqkit v2.13.0 (Shen et al., 2016), Tetranucleotide Frequency (TNF) using Biopython v1.81 (Cock et al., 2009) and relative synonymous codon usage (RSCU) using seqinr (translation table 11) (Charif et al., 2007). We obtained the differential of GC content (**Δ**GC) for each replicon in comparison with the largest replicon (chr1) from the corresponding genome. To quantify codon usage similarity, we calculated the Pearson correlation between the RSCU profiles of each replicon and the largest replicon (chr1) of the corresponding genome.

We used two complementary Hidden Markov Model (HMM)-based approaches to assess the presence of putative replicon-associated marker genes. The first one was the archaeal TIGRFAM marker set included in the GTDB Release 232 reference package. We used the 18 proteins included in the set to search against all predicted proteins using hmmscan from HMMER v3.4 (Mistry et al., 2013) with trusted cutoffs (--cut_tc). The second approach included archaeal marker protein families associated with chromosome maintenance (Cdc6/Orc1, MCM, RFC1, GINS15, GINS23 and RNA polymerase subunit Rpb2), partition systems (ParA and ParB families), and plasmid replication (RepA, RepB, RepC and Replitron HUH proteins). These markers are part of the Pfam HMM library distributed with the GTDB reference package. For each replicon, we calculated both the number of distinct marker families detected (marker presence), and the total number of proteins matching those marker families (marker hits). We incorporated these metrics into the replicon feature table and used them for downstream comparative analyses.

We evaluated the genomic features calculated for the archaeal replicons using dplyr (Wickham et al., 2026), tidyr (Wickham et al., 2025), ggplot2 (Wickham, 2016), vegan (Oksanen, 2017), and default R packages. For each genomic feature, we assessed its association with replicon length using Spearman’s rank correlation coefficient (ρ), and determined significance using a two-sided Spearman correlation test. Using a principal coordinates analysis (PCoA) based on Gower distances we visualized variation in genomic features among replicons. We assessed statistical differences among replicon types using permutational multivariate analysis of variance (PERMANOVA; adonis2, 9,999 permutations), and evaluated homogeneity of multivariate dispersions using betadisper (R package vegan). We assessed the associations between genomic features and the ordination axes using envfit with 999 permutations.

### Construction of protein families

To functionally annotate the 211 genomes, we first clustered all predicted proteins from the complete genome set using MMseqs2 release 13-45111 (Steinegger and Söding, 2017), using a minimum of 50% sequence identity and 80% sequence coverage. Clustering reduced computational redundancy by grouping similar proteins into representative protein families, which were subsequently used both for functional annotation and pangenome construction. Applying the same clustering framework across all 211 genomes provided a consistent definition of protein families for comparative analyses. We annotated the representative sequences from each cluster using eggNOG-mapper v2.1.13 (Huerta-Cepas et al., 2017) to assign Cluster of Orthologous Groups (COG) categories (Galperin et al., 2015). Throughout this study, protein families refer to MMseqs2 sequence clusters unless otherwise indicated. We generated the following tables for downstream analyses: (i) a protein-to-cluster assignment table, (ii) a protein metadata table containing genome, contig, and replicon information, (iii) an integrated protein-cluster table linking each protein to its genome, contig, and replicon, (iv) a table summarizing the abundance of each protein cluster across replicon categories, (v) a cluster-by-replicon abundance matrix, and (vi) a per-genome copy number table recording the number of copies of each protein cluster within each genome and replicon category. We used the latter to quantify gene family expansion, duplication frequency, and copy-number distributions. Additionally, to identify the proteins shared between the three different replicon units at phylum level, we determined shared protein clusters among chr1, chr2, and plasmids by constructing an UpSet diagram with the UpSetR package (Love et al., 2014).

### Protein family duplication and replicon gene sharing

We quantified protein family duplications for each replicon based on the protein clusters. For each genome–replicon combination, we determined the number of copies belonging to each protein family, and classified as duplicated those protein families represented by two or more copies within the same replicon. For each replicon, we calculated the total number of duplicated protein families, the total number of protein families, and the fraction of duplicated protein families. The relationship between replicon length and both the number and fraction of duplicated protein families was evaluated using Spearman’s rank correlation coefficient. Additionally, for each plasmid, we calculated the proportion of its protein families shared with chr1 of the same genome by dividing the number of shared protein families by the total number of plasmid protein families. We evaluated the association between plasmid length and the fraction of shared protein families with chr1 using Spearman’s rank correlation coefficient.

### Functional Annotation Analysis

We used the relative abundances of all COG functional categories to calculate Bray–Curtis dissimilarities among replicons. To calculate relative abundances, we accounted for the fact that some proteins were assigned to multiple COG categories (e.g. KL or MLT). In such cases, we counted each protein as contributing equally to each of its assigned categories, such that the total contribution of each protein sequence was one (i.e. a KL annotated protein would contribute 0.5 to category K and 0.5 to category L). We then constructed a PCoA to visualize differences in functional composition among replicon types, and assessed statistical significance using PERMANOVA (adonis2, 9,999 permutations), and homogeneity of multivariate dispersions using betadisper and envfit as previously described.

In addition to COG based functional annotation, we annotated proteins using FANTASIA (version 20241119, Singularity container), a protein language model-based annotation pipeline (Martínez-Redondo et al., 2025). Unlike homology-based methods, FANTASIA transforms protein sequences into high-dimensional embeddings using a protein language model and infers Gene Ontology (GO) terms as functional annotations based on similarity in the resulting embedding space. These results allowed us to annotate proteins lacking functional assignments from homology-based annotation (i.e. category S and non-annotated). We processed the resulting COG and GO terms tables using dplyr, tidyr, ggplot2, vegan, and R default packages.

### Halobacteriota pangenome construction

We also constructed three separate pangenomes using individual replicons as the units of comparison: a chr1 pangenome, a chr2 pangenome, and a plasmid pangenome. In each pangenome, protein clusters were classified as Core (present in ≥95% of replicons), Soft-core (present in ≥80%), Accessory (present in ≥15%), Rare (present in <15% but in more than one replicon), or Singleton (present in only one replicon). Thus, each protein family was assigned to a pangenome category according to the proportion of replicons within the corresponding pangenome in which it was detected. For example, for the chr1 pangenome, a protein cluster detected in 200 of 211 chr1 replicons would be classified as Core, whereas a cluster detected in 20 of 211 chr1 replicons would be classified as Accessory. Similarly, we constructed pangenomes at the family level using genomes instead of replicons as comparison units to obtain pangenomes with a higher biological interpretation. We constructed the family-level pangenomes only for families with ≥ 10 multipartite genomes (with either chr2 or plasmids) available (i.e. *Haladaptataceae* (n = 14), *Haloarculaceae* (n = 54), *Halobacteriaceae* (n = 20), *Haloferacaceae* (n = 58), and *Natrialbaceae* (n = 44)).

### Shared functions among replicons in Halobacteriota families with secondary chromosomes

To investigate protein-family sharing among replicons within individual genomes, we selected genomes containing at least one chr2. We selected a total of nine genomes: *Haladaptatus* sp020618475 (GCF_020618475.1), *Halocatena* (GCF_027563145.1), *KZCA124* sp017357405 (GCF_017357405.2), *Haloarcula* (GCF_051122755.1), *Haloprofundus* sp020150815 (GCF_020150815.1), *Halorubrum lacusprofundi* (GCF_000022205.1), *Haloarcula marismortui* (GCF_000011085.1), and *Haloarcula hispanica* (GCF_000223905.1 and GCF_000504565.1). For each genome, we constructed a network in which nodes represented replicons and edges represented the number of protein families shared between pairs of replicons. For network visualization we used igraph (Csárdi and Nepusz, 2006) and ggraph (Pedersen, 2025) R packages. Then, we classified shared protein families according to the combination of replicon types in which they occurred (e.g., chr1–plasmid, chr2–plasmid, chr1–chr2, or shared among all replicons). Finally, we evaluated composition similarity by comparing the GC content of every gene belonging to a shared protein family with the GC content of the replicon carrying that gene. We calculated the absolute GC-content difference (|ΔGC|) for each gene and summarized these values as the mean |ΔGC| for each genome and shared-cluster category.

## RESULTS

### Genomic landscape of chromosomes, secondary chromosomes, and plasmids

We downloaded all 626 publicly available complete archaeal genomes from RefSeq that carried at least one additional replicon besides chr1, irrespective of replicon length. After taxonomic annotation, we selected only Halobacteriota genomes, comprising a total of 211genomes (33.7% of all complete archaeal genomes with at least one additional replicon), distributed across ten families (Table S1). These genomes contained a total of 766 replicons, classified into three types: 211 chr1, 9 chr2, and 546 plasmids. Replicon sizes ranged from 0.2 to 4.8 Mb for chr1, from 0.28 to 2.0 Mb for ch2, and from 0.001 to 1.6 Mb for plasmids (Table S1).

To characterize and compare the three types of replicons, we analysed a range of genomic features: length, codon usage, GC content, tetranucleotide frequencies (TNF) values, number of chr1 markers hits, number of GTDB markers hits, number of partition markers hits, and number of plasmid markers hits. The results showed that Halobacteriota secondary replicons, that is, replicons classified as chr2 or plasmids, form a continuum along codon usage, GC content, length, and TNF (Fig. 1A, Fig. S1A). Interestingly, chr1 is also distributed along this continuum, although they occupy a more restricted region of the distribution (Fig. 1A, Fig. S1A). A similar pattern can be appreciated in the number of plasmid hits, and to a lesser extent in partition and chr1 hits. These markers provide information about replicon maintenance and identity: plasmid replication proteins support plasmid-like replication, whereas partition and chr1-associated markers indicate mechanisms involved in replicon segregation and chromosome maintenance, respectively. The feature that most clearly distinguishes chr1 from secondary replicons is the GTDB markers category, which effectively discriminates chr1 from all other replicon types. Chr2 tends to occupy an intermediate position between plasmids and chr1, particularly with respect to length, number of chr1 hits, and, to a lesser extent, TNF and codon usage patterns (Fig. 1A). Longer secondary replicons tend to exhibit more chromosome-like properties, showing positive associations with multiple chromosome-related features, including chromosome marker genes, partition genes, codon usage similarity to chr1, and TNF similarity. In contrast, GC content and GTDB marker abundance showed comparatively weak associations with replicon size (Fig. S1B).

**Figure 1.**
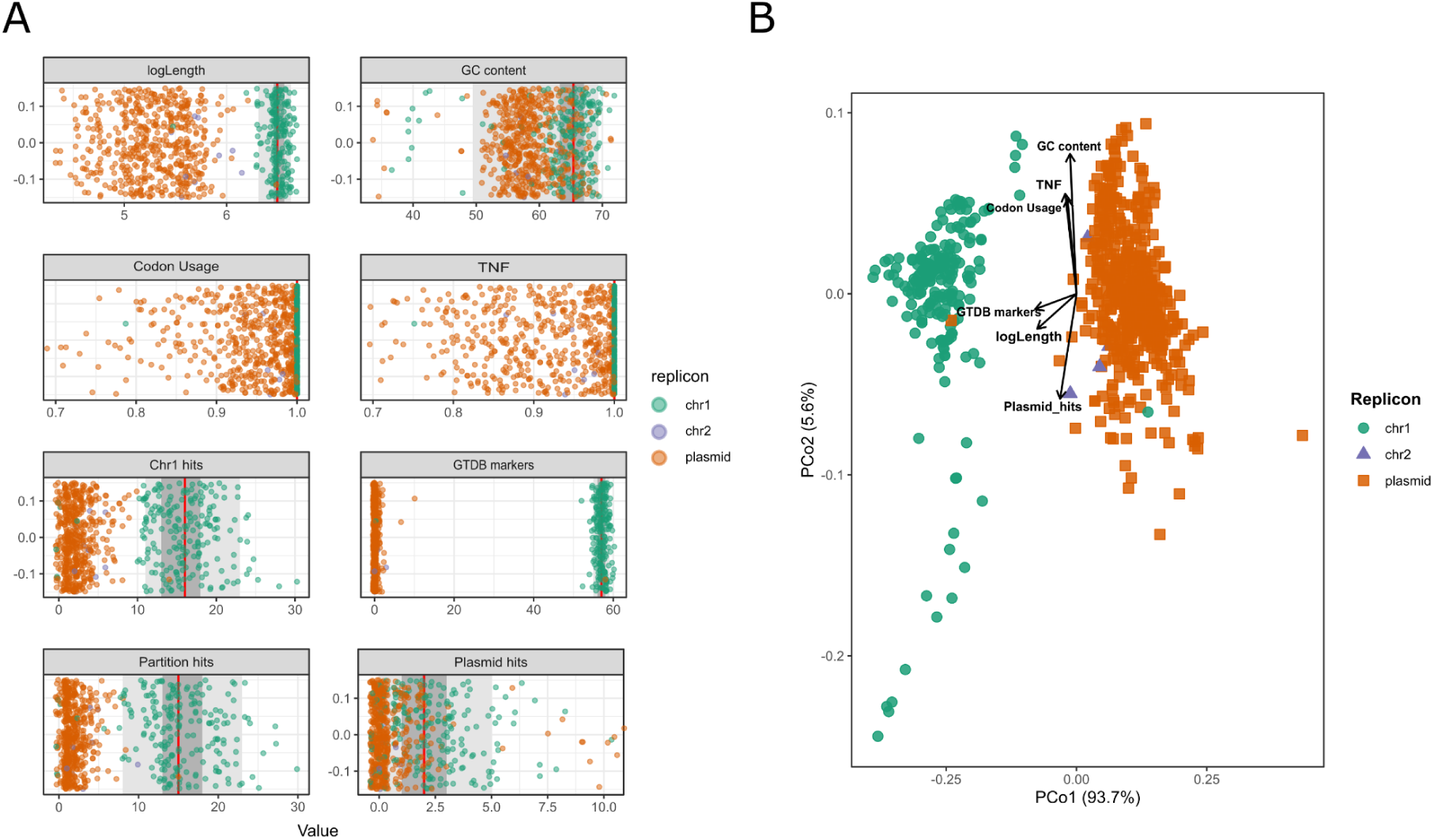
Genomic landscape of Halobacteriota replicons. (A) Distribution of all replicons in the dataset according to their value in each of the evaluated features. Each x-axis corresponds to the range of values associated with each of the features. Outliers in Codon Usage and TNF features were removed for better visualization. Graphs including outliers are available in Supplementary Material (Fig. S1). (B) PCoA ordination showing all replicons in the Halobacteriota dataset. Each point corresponds to a replicon and shape and color indicate replicon type. Vectors corresponding to the different genomic features are shown. Statistical significance was evaluated through PERMANOVA, R² = 0.853, F = 2117.51, P < 0.0001, and PERMDISP, F = 6.58, P = 0.006.

To determine whether these genomic features collectively define replicon identity, we projected all replicons into a multivariate feature space using Principal Coordinate Analysis (Fig. 1B, Fig. S1C). The first coordinate, explaining 93.7% of the variance, primarily separates chromosomes from plasmids, while chr2 replicons occupy an intermediate region, consistent with a genomic continuum rather than discrete replicon classes. The three replicons differed significantly (PERMANOVA, R² = 0.853, F = 2117.51, P < 0.0001), although differences in multivariate dispersion were also detected (PERMDISP, F = 6.58, P = 0.006). PCoA1 was primarily associated with replicon length and chromosome-associated features, including GTDB markers, chromosome marker genes, and partition genes (all P < 0.001). In contrast, PCoA2 (5.6% of the variance) was primarily associated with sequence composition, showing positive correlations with GC content, tetranucleotide frequency similarity, and codon usage similarity, and a negative correlation with plasmid marker abundance (all P < 0.001; Fig. 1B). Within the chr1 cluster, 16 chromosomes form a more dispersed group in the bottom left region of the ordination. These chromosomes correspond to all the archaeal genomes from the *Methanosarcinaceae* family and some members of the *Methanotrichaceae* (*Methanothrix soehngenii*), *Haloferacaceae* (*Haloquadratum walsbyi*), and *Archaeoglobaceae* (*Archaeoglobus_B profundus*) families (Fig. S1C). These chr1 have a mean GC content of 42.31%, lower than that of the chr1 replicons located in the upper group (65.66%). The remaining genomic features do not differ between the upper and lower groups (Table S2).

Based on the eight selected genomic features, secondary replicons do not appear to fall into distinct biological categories, but rather form a genomic continuum from plasmid-like to chromosome-like states. Increasing replicon size is associated with progressively more chromosome-like features, supporting a gradual transition in replicon identity and organization.

### Distribution of functional content among Halobacteriota replicons

Next, we examined whether the three replicon types exhibited distinct functional profiles. Functional annotation using the COG database showed that after excluding the function unknown category (S), the most abundant categories were transcription (K), amino acid metabolism (E), replication and repair (L), inorganic ion transport and metabolism (P), carbohydrate metabolism (G), and signal transduction (T). Although the distribution of functional categories was broadly similar across the three replicon types (Table 1), COG functional composition differed significantly among them (PERMANOVA, R² = 0.147, F = 64.86, P < 0.0001). Multivariate dispersion also differed among replicon types (PERMDISP, F = 428.34, P = 0.001), indicating differences in within-group variability (Fig. 2A). Specifically, genes involved in inorganic ion metabolism and carbohydrate metabolism (P and G) are more abundant in chr2, while signal transduction (T) is less abundant in chr2 than in chr1 and plasmids (Table 1). The most abundant COG categories comprised a broad range of functions and were not driven by a small number of highly abundant proteins (Table S3).

**Figure 2.**
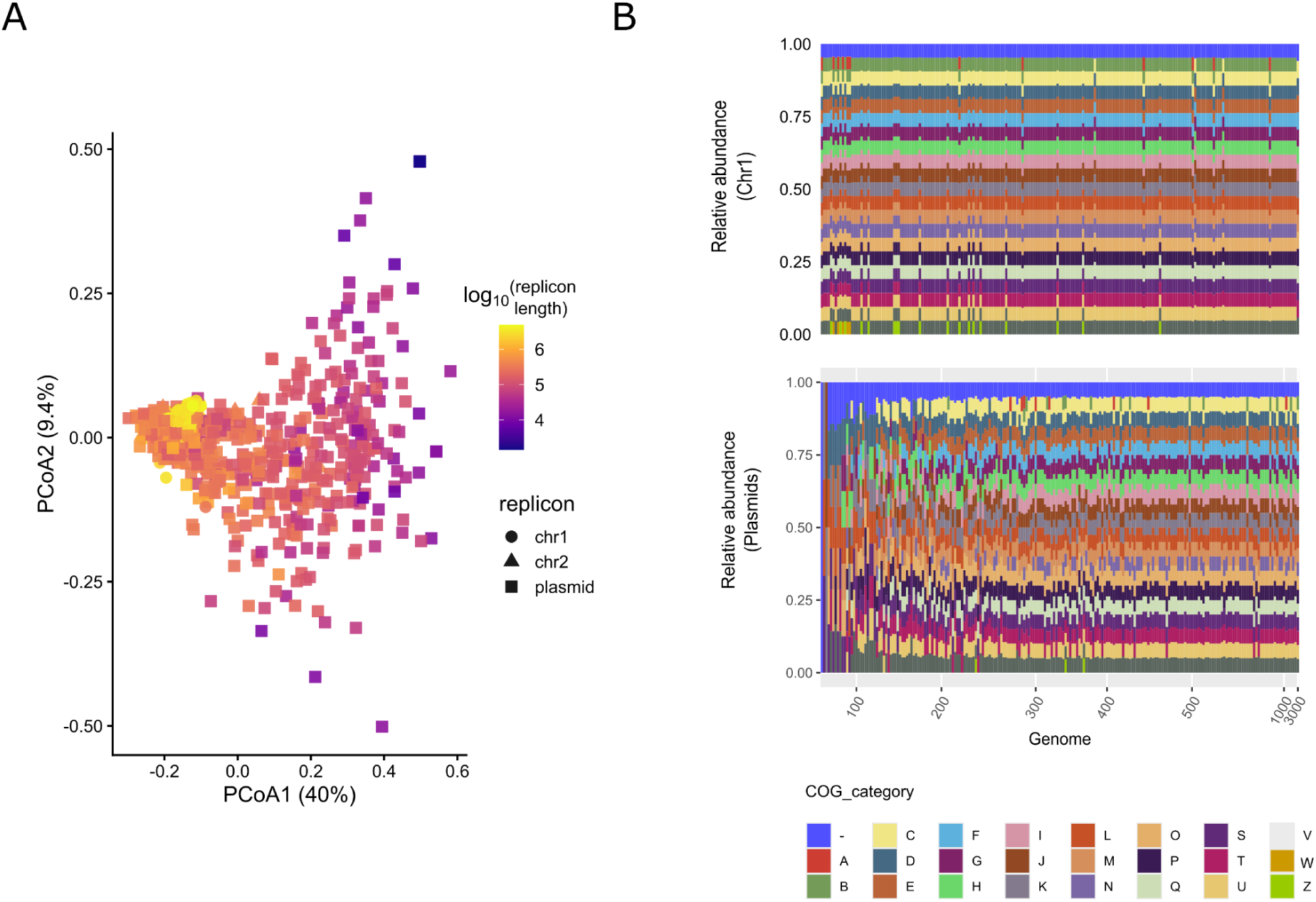
Changes in functional content and duplications according to replicon size. (A) PCoA ordination where each point represents a replicon of the Halobacteriota dataset. Color corresponds to replicon size and shape replicon type. Statistical significance was evaluated through PERMANOVA, R² = 0.147, F = 64.86, P < 0.0001 and PERMDISP, F = 428.34, P = 0.001. (B) Barplots showing COG functional content per replicon. The lower plot shows secondary replicons ordered by increasing length, with the corresponding chr1 replicons shown in the upper barplot according to the ordering of their associated secondary replicons. COG functional content of the lower plot becomes remarkably similar to the pattern in the upper plot as replicon length increases.

**Table 1.**
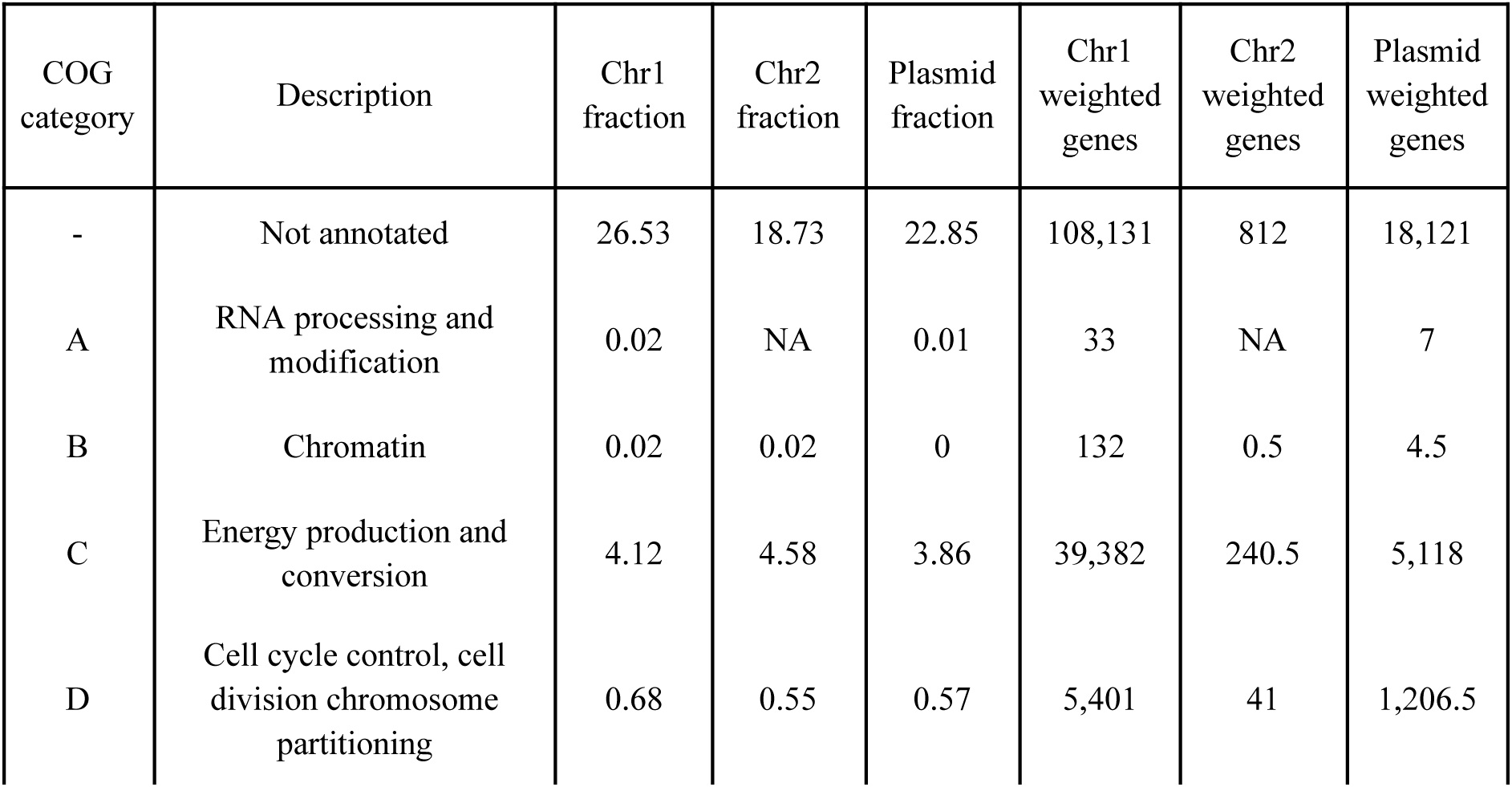
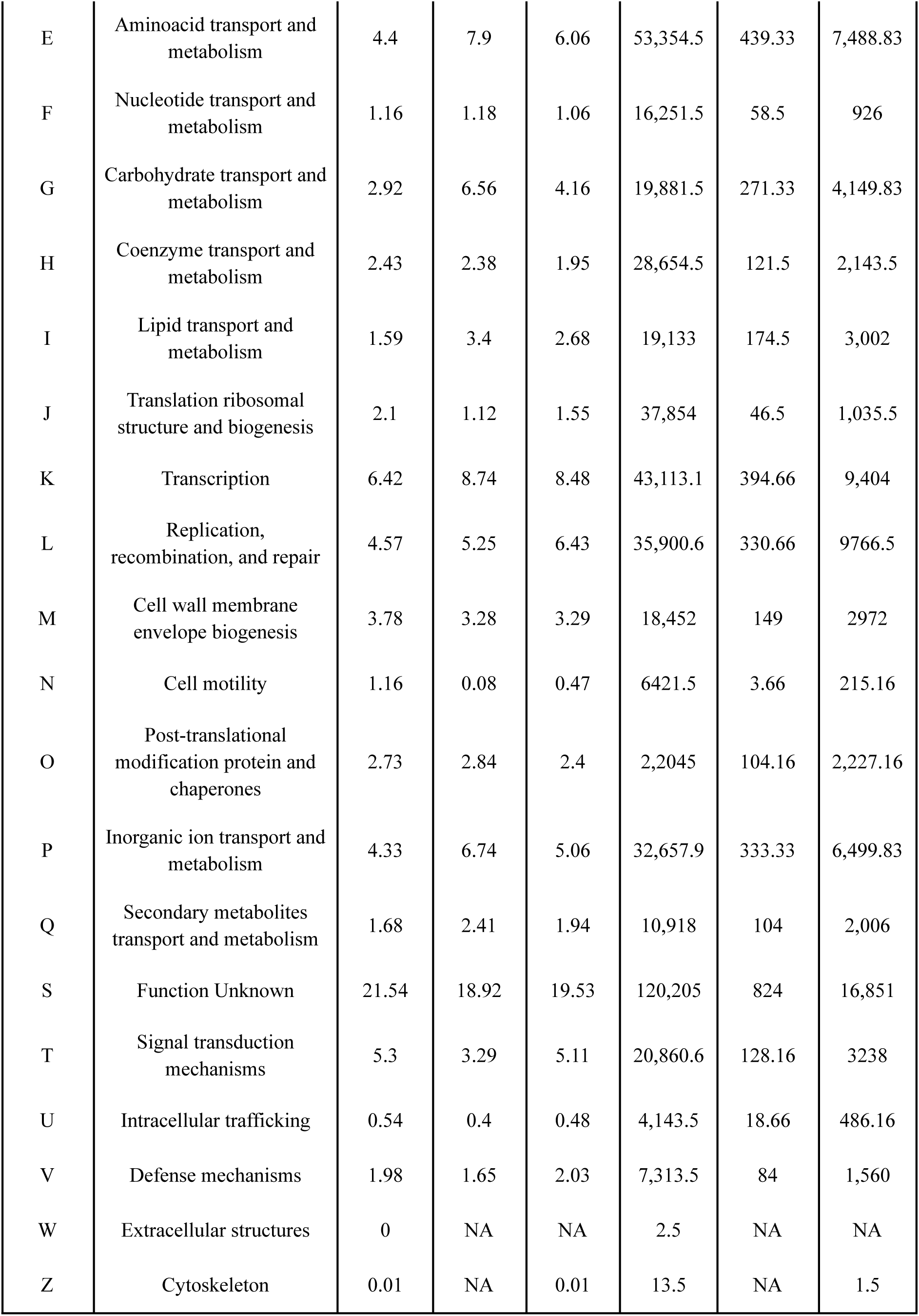
COG category composition across replicon types. Relative abundance (%) of distinct gene clusters assigned to each COG functional category in the primary chromosome (Chr1), secondary chromosomes and additional replicons (Chr2+), and plasmids. Weighted protein counts are also shown for each category; proteins assigned to multiple COG categories contribute proportionally to each category.

| COG category | Description | Chr1 fraction | Chr2 fraction | Plasmid fraction | Chr1 weighted genes | Chr2 weighted genes | Plasmid weighted genes |
| --- | --- | --- | --- | --- | --- | --- | --- |
| - | Not annotated | 26.53 | 18.73 | 22.85 | 108,131 | 812 | 18,121 |
| A | RNA processing and modification | 0.02 | NA | 0.01 | 33 | NA | 7 |
| B | Chromatin | 0.02 | 0.02 | 0 | 132 | 0.5 | 4.5 |
| C | Energy production and conversion | 4.12 | 4.58 | 3.86 | 39,382 | 240.5 | 5,118 |
| D | Cell cycle control, cell division chromosome partitioning | 0.68 | 0.55 | 0.57 | 5,401 | 41 | 1,206.5 |
| E | Aminoacid transport and metabolism | 4.4 | 7.9 | 6.06 | 53,354.5 | 439.33 | 7,488.83 |
| F | Nucleotide transport and metabolism | 1.16 | 1.18 | 1.06 | 16,251.5 | 58.5 | 926 |
| G | Carbohydrate transport and metabolism | 2.92 | 6.56 | 4.16 | 19,881.5 | 271.33 | 4,149.83 |
| H | Coenzyme transport and metabolism | 2.43 | 2.38 | 1.95 | 28,654.5 | 121.5 | 2,143.5 |
| I | Lipid transport and metabolism | 1.59 | 3.4 | 2.68 | 19,133 | 174.5 | 3,002 |
| J | Translation ribosomal structure and biogenesis | 2.1 | 1.12 | 1.55 | 37,854 | 46.5 | 1,035.5 |
| K | Transcription | 6.42 | 8.74 | 8.48 | 43,113.1 | 394.66 | 9,404 |
| L | Replication, recombination, and repair | 4.57 | 5.25 | 6.43 | 35,900.6 | 330.66 | 9766.5 |
| M | Cell wall membrane envelope biogenesis | 3.78 | 3.28 | 3.29 | 18,452 | 149 | 2972 |
| N | Cell motility | 1.16 | 0.08 | 0.47 | 6421.5 | 3.66 | 215.16 |
| O | Post-translational modification protein and chaperones | 2.73 | 2.84 | 2.4 | 2,2045 | 104.16 | 2,227.16 |
| P | Inorganic ion transport and metabolism | 4.33 | 6.74 | 5.06 | 32,657.9 | 333.33 | 6,499.83 |
| Q | Secondary metabolites transport and metabolism | 1.68 | 2.41 | 1.94 | 10,918 | 104 | 2,006 |
| S | Function Unknown | 21.54 | 18.92 | 19.53 | 120,205 | 824 | 16,851 |
| T | Signal transduction mechanisms | 5.3 | 3.29 | 5.11 | 20,860.6 | 128.16 | 3238 |
| U | Intracellular trafficking | 0.54 | 0.4 | 0.48 | 4,143.5 | 18.66 | 486.16 |
| V | Defense mechanisms | 1.98 | 1.65 | 2.03 | 7,313.5 | 84 | 1,560 |
| W | Extracellular structures | 0 | NA | NA | 2.5 | NA | NA |
| Z | Cytoskeleton | 0.01 | NA | 0.01 | 13.5 | NA | 1.5 |

We used FANTASIA to infer the functions of proteins assigned to COG category S (Function unknown) as well as proteins lacking functional annotation, but found no differences among the three replicons (Table S4). Across all three replicon types, the majority of these proteins were associated with cellular components related to cytoplasmic membrane, integral components of the membrane, and cytoplasmic cellular functions (Fig. S2). In the molecular function category, DNA binding and metal ion binding were the most prevalent FANTASIA annotations across all replicon types.

Overall, COG functional annotation revealed a broadly similar distribution of functions across replicon types, with no clear functional partitioning of proteins involved in either core cellular processes or environmental sensing and nutrient acquisition.

### Secondary replicons become progressively more chromosome-like with increasing length

Although the distribution of functional categories was similar across all three types of replicons, we identified an association between length and the functional content of secondary replicons. Variation in COG functional composition follows a gradient of replicon size, with length being strongly associated with the first PCoA axis (R² = 0.562, P = 0.001) (Fig. 2A). Across Halobacteriota genomes, chr1 exhibited a remarkably conserved functional composition, with the relative abundance of COG categories remaining largely constant despite considerable variation in genome size (Fig. 2B). In contrast, secondary replicons displayed substantially greater functional heterogeneity, particularly among smaller replicons. However, as secondary replicon length increased, their functional profiles became progressively more stable and increasingly converged toward those of chr1 (Fig. 2B). Categories related to aminoacid transport and metabolism (E), carbohydrate transport and metabolism (G), energy production and conversion (C), coenzyme transport and metabolism (H), and secondary metabolite biosynthesis (Q) were positively correlated with replicon length, whereas non-annotated proteins and replication and repair proteins (L) showed a strong negative correlation (ρ = -0.59, FDR < 0.001 and ρ = -0.36, FDR < 0.001, respectively, Fig. S3A). A similar trend was observed along the TNF gradient, although it was less pronounced than the trend associated with replicon length. As the TNF of secondary replicons became increasingly similar to that of the chr1, the functional composition of plasmids also converged toward that of the chromosome at the COG category level (Fig. S3B). Proteins involved in energy production and conversion (C), aminoacid metabolism (E), and carbohydrate transport and metabolism (G) were positively correlated with TNF, and replication and repair (L) was negatively correlated (Fig. S3C).

Consistent with these results, the fraction of plasmid protein families shared with the host chr1 increased significantly with plasmid length (Spearman’s ρ = 0.278, P < 0.001; Fig. 3A), indicating greater protein-family overlap with chr1 in larger plasmids. We next examined whether gene-family duplication was similarly associated with replicon size. Replicon length was strongly correlated with the number of duplicated protein families (ρ = 0.926, P < 0.001) (Fig. 3B), and this relationship remained significant when chr1 (ρ = 0.838, P < 0.001), chr2 (ρ = 0.917, P = 0.001), and plasmids (ρ = 0.820, P < 0.001) were analyzed separately. Together, these results show that increasing replicon size is associated with greater gene-family duplication across replicon types and, in plasmids, greater protein-family overlap with the primary chromosome (Fig. 3A).

**Figure 3.**
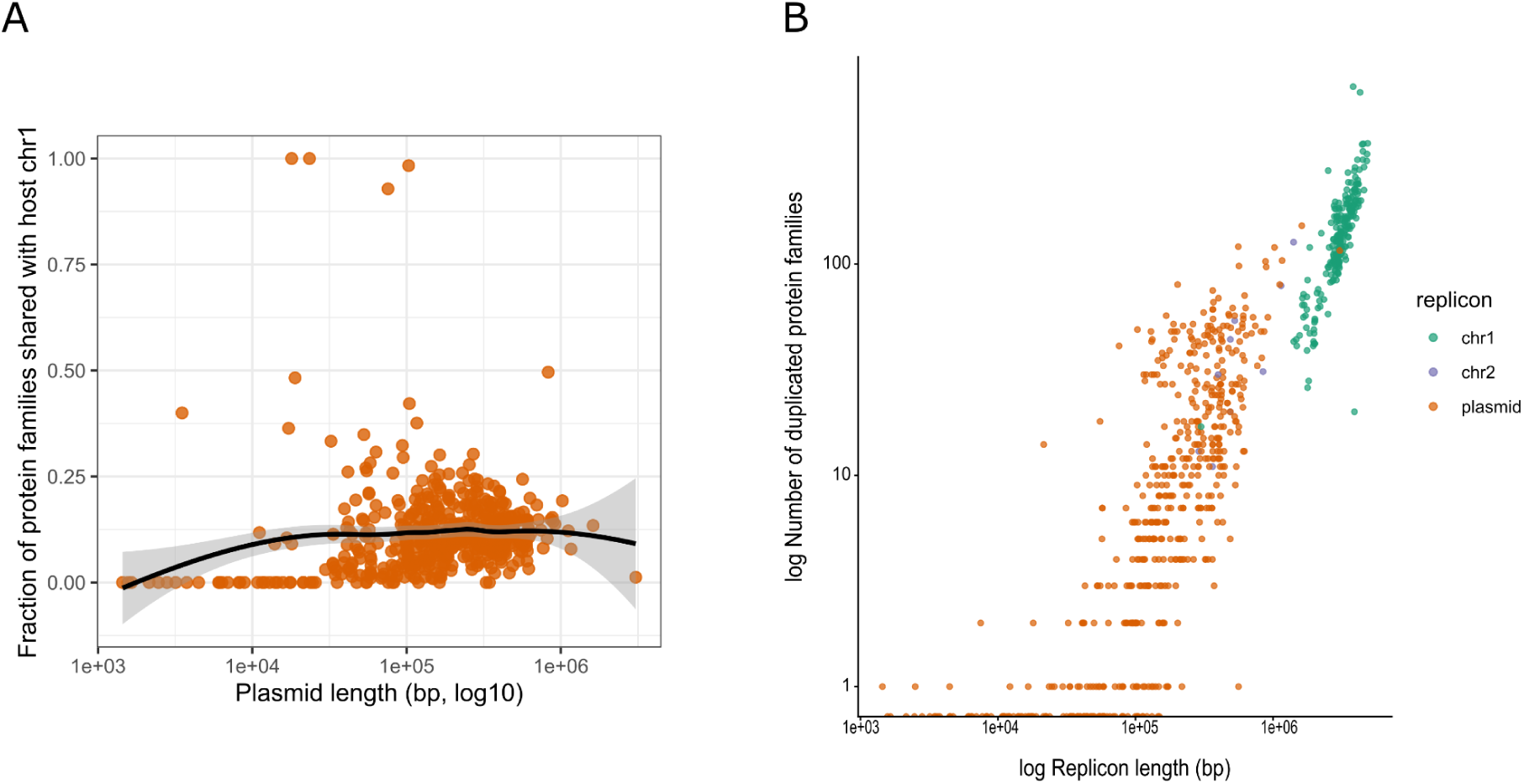
Chromosome-like features increase with replicon size. (A) Dispersion plot showing the number of shared proteins between secondary replicons and chr1 as secondary replicon length increases. Each point represents a secondary replicon and the black line represents a LOESS-smoothed trend, with the shaded area indicating the 95% confidence interval. (B) Dispersion plot representing the number of duplicated protein families in each replicon according to their length. Each point represents a replicon and colors indicate replicon type. Both axes are in log_10_ scale.

Overall, these patterns further support a continuum in secondary-replicon organization, in which larger plasmids show greater functional overlap with chr1 and increased gene-family duplication. Thus, the shift toward chromosome-like properties with increasing replicon size extends beyond sequence composition to protein-family content and organization.

### Shared proteins among Halobacteriota replicons and pangenome construction

To evaluate the degree of functional integration of secondary replicons, we conducted several analyses to determine the fraction of protein families shared among replicon types. Our first analysis identified a small core comprising only 2,1% of all protein families: of the 122,894 protein-family clusters in the dataset 2,593 were shared among chr1, chr2, and plasmids, constituting the Phylum Core (Fig. S4A). The Phylum Core was mainly dominated by proteins involved in aminoacid metabolism, such as aminoacid transporters and enzymes involved in branched-chain aminoacid metabolism (COG E, n = 218), transcriptional regulators and DNA binding proteins (COG K, n = 209), and inorganic ion transport and metabolism, including sugar, phosphorus, nickel, cobalt, and other heavy metal transporters (COG P, n = 187) (Table S5).

At phylum level, chr2 and chr1 shared only 180 protein families (Fig. S4A), predominantly associated with environmental sensing and regulation (e.g., transcriptional regulators, PAS/PAC domain proteins, histidine kinases), transport systems (ABC and major facilitator superfamily transporters), carbohydrate metabolism (e.g., transketolases, chitinases and pectin methylesterases), and genome maintenance (e.g., integrases and helicases), whereas core functions were largely absent in this shared set. Similarly, only 514 clusters were shared between chr2 and plasmids (Fig. S4A), mainly associated with environmental sensing and regulation (e.g., transcriptional regulators, PAS/PAC proteins and histidine kinases), carbohydrate metabolism (e.g., β-galactosidases, trehalose utilization proteins and xylose isomerases), genome maintenance and mobility (e.g., ParA ATPases, integrases, transposases and Type IV secretion proteins), transporters (ABC, MFS and Na++/H++ antiporters), and stress-response proteins (e.g., UspA and F420420-dependent oxidoreductases) (Table S5). Lastly, 13,414 protein families were shared between plasmids and chr1 (Fig. S4A). These were predominantly associated with energy metabolism and electron transfer, as well as with components of respiratory complexes and ATP synthase, indicating that conserved metabolic and bioenergetic functions constitute a major fraction of the shared accessory gene repertoire (Table S5). An important limitation of this analysis is the considerably low number of chr2 replicons (n = 9) compared with chr1 (n = 211) and plasmids (n = 546).

To obtain a higher resolution than that obtained by identifying shared protein families across chr1, chr2, and plasmids, we constructed pangenomes. We followed two different approaches to construct pangenomes: (i) using replicons as comparison units, and (ii) using genomes as comparison units. When using replicons, we obtained three pangenomes, one per each type of replicon. Across all three replicon types, singletons constituted the largest pangenome category, comprising 55.6% of all protein families in the chr1 pangenome to up to 81.8% in the chr2 pangenome (Fig. S4B). Consistent with this high level of diversity, we were unable to identify any proteins that met the criteria to be classified as core protein in either the chr2 or plasmid pangenomes (Fig. S4B). Only one single protein family comprised the soft-core of the chr2 pangenome (ATPase associated with multiple activities, Table S6). In contrast, chr1 pangenome contained 118 protein families in the core genome, and 730 protein families in the soft-core genome. As expected, chr1 core protein families are involved in core cellular processes, including translation, DNA replication and repair, transcription, and central metabolism (Table S6). In contrast, plasmids were characterized by a predominance of rare and singleton protein families (99.9% of proteins) and a remarkably low number of accessory protein families (n = 35), despite the high number of plasmids present in the dataset (n = 546) (Fig. S4B).

Next, we computed one pangenome per Halobacteriota family using genomes as the comparison units (approach ii). As observed in the previous analysis, singleton protein families constituted approximately half of the pangenome across all five families, whereas core and soft-core families together represented less than 10% (Table 2). Differences among families were primarily driven by the relative proportions of accessory and rare protein families, with *Halobacteriaceae* and *Haladaptataceae* showing higher proportion of accessory families, whereas *Haloarculaceae*, *Haloferacaceae*, and *Natrialbaceae* exhibiting a greater proportion of rare families. The distribution of pangenome categories (i.e. core, soft-core, accessory, rare, and singleton) were not partitioned among replicon types. Instead, proteins from all categories were distributed across all three replicons (Fig. 4A). Core proteins were not exclusively located on chr1, nor were singletons restricted to plasmids. Chr1 contained the majority of proteins overall and also harboured the largest number of proteins in each pangenome category, including core, soft-core, accessory, rare, and singleton proteins (Fig. 4A). This pattern was consistent across all five families analyzed, with chr1 containing the largest number of proteins in each category, followed by plasmids and chr2.

**Figure 4.**
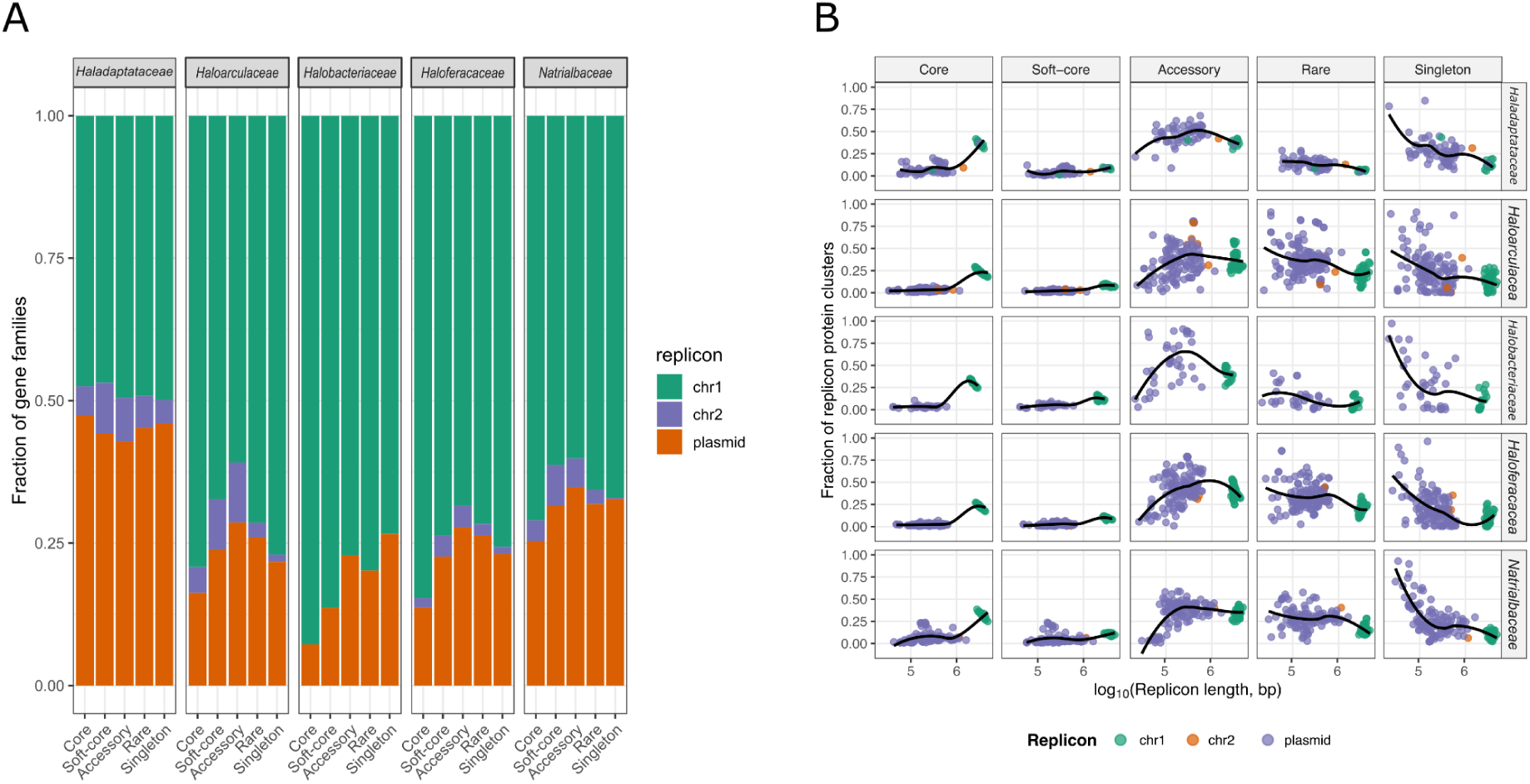
Pangenomes constructed at a family taxonomic level. (A) Distribution of protein families from all pangenome categories among the different replicon types. (B) Dispersion plots depicting the number of protein families of each category per each individual replicon according to replicon length. Each point represents a replicon, color indicates replicon type. A LOESS-smoothed trend is shown for each plot. Both panels show the plots for *Haladaptataceae*, *Haloarculaceae*, *Halobacteriaceae*, *Haloferacaceae*, and *Natrialbaceae* families.

**Table 2.** Distribution of protein families across core, soft-core, accessory, rare, and singleton categories in family-level pangenomes.

| Family | Core | Soft-core | Accessory | Rare | Singleton | Total |
| --- | --- | --- | --- | --- | --- | --- |
| *Haladaptataceae* | 1,184<br>(6.26%) | 344<br>(1.82%) | 4,619<br>(24.42%) | 2,482<br>(13.12%) | 10,285<br>(54.38%) | 18914 |
| *Haloarculaceae* | 765<br>(1.76%) | 319<br>(0.73%) | 4,166<br>(9.57%) | 14,695<br>(33.76%) | 23,580<br>(54.18%) | 43525 |
| *Halobacteriaceae* | 702<br>(6.46%) | 348<br>(3.20%) | 3,038<br>(27.95%) | 1,628<br>(14.98%) | 5,155<br>(47.42%) | 10871 |
| *Haloferacaceae* | 692<br>(1.99%) | 332<br>(0.95%) | 4,412<br>(12.68%) | 11,961<br>(34.37%) | 17,400<br>(50.00%) | 34797 |
| *Natrialbaceae* | 1,089<br>(3.14%) | 439<br>(1.27%) | 3,944<br>(11.38%) | 9,788<br>(28.25%) | 19,389<br>(55.96%) | 34649 |

Across all five Halobacteriota families, replicon size was strongly associated with pangenome composition. Larger replicons contained progressively a higher proportion of core and soft-core protein families, whereas the fraction of singleton families declined consistently with increasing replicon length (Fig. 4B). Accessory protein families exhibited a peak in intermediate-sized replicons and decreased in the largest replicons. Similarly, rare protein families also tended to decline with replicon size.

Overall, Halobacteriota replicons displayed highly heterogeneous gene repertoires, particularly among smaller secondary replicons. However, this diversity decreased with increasing replicon size: larger replicons contained greater proportions of conserved core and soft-core protein families, whereas smaller replicons were dominated by rare and lineage-specific genes.

### Some protein families are shared among all replicons within a genome

Given that all replicon types shared similar COG functional profiles across the entire Halobacteriota dataset (Table 1), we sought to investigate functional conservation in more detail. To this end, we evaluated shared protein families within a subset of nine genomes selected on the basis of harboring a chr2 replicon, regardless of the number of plasmids contained. Shared protein families between the replicons were individually analyzed for each of the nine Halobacteriota selected genomes. In each genome, all replicons shared protein families, with *Haladaptatus* sp sharing the highest number of shared protein clusters (n = 717), followed by *KZCA124* sp (n = 203), and *Haloprofundus* sp being the genome with fewest shared protein families (n = 55) (Table S7). In all cases, chr1 shared protein clusters with virtually every other chr2 and most plasmids in the genome (Fig. S5). However, network topology varied markedly among genomes, ranging from simple chr1-chr2 systems to highly interconnected multipartite architectures involving multiple plasmids. Similarly, the distribution of shared protein-family classes differed substantially among species. For example, shared protein families in *Halocatena* and *KZCA124* sp were more abundant between chr1 and chr2 than with plasmids, whereas *Haladaptatus* sp exhibited extensive protein family sharing among chromosomes and plasmids, including many protein families shared only among plasmids or across all replicon types (Fig. S5).

Functional annotation of the genomes from these nine Halobacteriota families revealed that chr1-chr2 shared clusters were predominantly associated with housekeeping functions, including translation, transcription, DNA replication and central metabolism, whereas chr1-plasmid and plasmid-plasmid shared clusters were enriched in regulatory proteins and mobile genetic elements such as integrases and transposases. Transporter-related proteins, however, were shared across all three replicon types (Table S7). Protein families lacking COG annotations were re-annotated using the protein language model-based FANTASIA pipeline. Besides recovering broad functional categories such as DNA binding and membrane-associated proteins, FANTASIA revealed genome-specific functional signatures (Table S8). For example, in *Haladaptatus* sp, proteins shared between chr1 and plasmids were predominantly associated with membrane-associated proteins, gas vesicle formation and DNA-binding functions, suggesting roles in environmental interaction, whereas proteins shared between chr1 and chr2 were mainly associated with central metabolic functions. In contrast, *KZCA124* sp showed an enrichment of secretion-related proteins shared between chr1 and plasmids, including Type IV secretion system components, together with membrane-associated proteins, consistent with a greater contribution of mobile genetic element-associated functions to the shared accessory genome. Likewise, FANTASIA inferred function for unannotated proteins from *Haloprofundus* sp, linking them to transmembrane transport and components of the archaeal transcription machinery (TFIIH). Together, these observations indicate that multipartite archaeal genomes do not conform to a single organizational model. Instead, they exhibit lineage-specific patterns of protein-family partitioning and functional integration across replicons.

The tendency of larger secondary chromosomes to resemble chr1 was also evident in the analysis of protein families shared among replicons across the nine Halobacteriota families examined (Fig. S5, Table S7). Despite the functional differences described above, genes belonging to shared protein families generally showed small to moderate GC-content differences relative to their host replicons (mean |ΔGC| ≈ 2–6%; Fig. S6), although the magnitude varied among genomes and categories of shared families. Families shared exclusively among plasmids or between plasmids and chromosomes occasionally showed larger compositional differences, with the highest |ΔGC| observed for families shared among all plasmids in the genome of *Haloarcula marismortui* (Fig. S6).

Altogether, our results show that the distribution of shared protein families across replicons varied among genomes, revealing lineage-specific patterns of functional integration. Despite this variability, chr1–chr2 sharing was generally enriched in conserved cellular functions, whereas plasmid-associated sharing involved more regulatory, transport, and mobile-element functions, with larger secondary replicons showing greater similarity to chr1.

## DISCUSSION

### Halobacteriota replicons are distributed along a structural continuum and a compositional axis

Halobacteriota thrive in highly saline environments, including near-saturation conditions (Becker et al., 2014), and many possess multipartite genomes containing one or more secondary replicons. The prevalence of multipartite genome architectures makes Halobacteriota a suitable system for investigating the genomic characteristics and evolutionary relationships of secondary replicons. We therefore evaluated genomes containing one or more replicons in addition to chr1, regardless of replicon size, using a range of genomic features reflecting distinct evolutionary constraints. Replicon length is related to gene acquisition history, GC content reflects long-term mutational equilibrium, codon usage indicates the degree of adaptation of the replicon to its host, and TNF reflects local sequence composition. Finally, partition, chr1, and plasmid markers were used to assess the abundance of replication and segregation functions associated with each replication type.

Contrary to our expectations, we found that the replicons were distributed along a continuum rather than forming discrete categories (Fig. 1A). The GTDB markers were the only evaluated feature that consistently distinguished chr1 from the other replicon types, underscoring their value for identifying primary chromosomes in archaeal genomes and supporting their use for taxonomic classification. Similarly, partition systems (e.g., ParA/ParB, included in our partition markers set) were also enriched in chr1, as expected given their typical association with large, low-copy number replicons, where they ensure accurate segregation during cell division (Baxter and Funnell, 2014). Core functions and partition systems, such as those included in our marker sets, may be less prone to horizontal transfer because they are essential and are highly integrated into the cellular networks (Cohen et al., 2011), explaining their enrichment in chr1. However, core functions have also been reported in plasmids (Ng et al., 1998; Gophna and Altman-Price, 2022), suggesting transfer between chr1 and plasmids. This raises the possibility that similar gene transfer events may also occur between chr 1 and secondary replicons in archaeal genomes. Indeed, positive hits to plasmid replication proteins (RepA, RepB, RepC and Replitron HUH proteins) were detected not only in plasmids but also consistently in chromosomes (Fig. 1A). The presence of plasmid-like features in archaeal chromosomes suggests that plasmid may have integrated into chr1 during their evolutionary history (Hawkins et al., 2013).

GC content showed a gradual distribution, consistent with a slow amelioration toward a more chromosome-like GC composition over evolutionary time. In contrast, TNF and codon usage patterns in secondary replicons showed a less gradual distribution, with values clustering more closely to chr1, suggesting that translation optimization evolves relatively rapidly, or at least at a faster rate than GC content.

Altogether, secondary replicon evolution appears to follow a continuum shaped by two classes of genomic features: threshold traits (i.e. chr1 and partition markers, GTDB classification, and replicon length) and amelioration traits (i.e. GC content, TNF, codon usage) traits. Threshold traits may reflect evolutionary tipping points at which secondary replicons transition to stably inherited and functionally integrated components of the genome. On the other hand, amelioration traits reflect the progressive adaptation of sequence composition to the host genome through mutation and selection, resulting in a continuum of evolutionary states rather than discrete replicon categories. Importantly, these compositional changes may start before a secondary replicon acquires a chromosomal-like functional identity. As a replicon becomes sufficiently integrated and essential to the host genome, selective pressures may favour its transition toward chromosome-like maintenance and inheritance. This transition could eventually lead to secondary replicons crossing an evolutionary “threshold” and joining the chr1-like replicons characterized by high numbers of partition systems and chromosome marker genes (Fig. 1A). The mechanisms underlying the acquisition of these genes are beyond the scope of this study but warrant further investigation.

Consistent with the patterns observed for individual genomic features, the PCoA shows a clear separation between threshold and amelioration traits. Vectors associated with amelioration traits (GC content, TNF and codon usage) pointed in a similar direction, whereas threshold traits (GTDB markers, chromosome hits and partition systems) primarily defined the axis of chromosome-like integration (Fig. 1B). This pattern suggests that amelioration traits reflect compositional similarity between the secondary replicons and the archaeal host genome, indicative of their shared evolutionary relationship, whereas threshold traits capture the extent to which secondary replicons have become integrated into chromosome-like maintenance and inheritance processes. The weak clustering of genomes by archaeal family further suggests that these patterns are driven mainly by the evolutionary state of the replicon rather than by host phylogeny. (Fig. S1C).

### Functional repertoires become progressively homogenized as replicons become chromosome-like

Functions were evenly distributed among the three replicon types at the phylum level (Table 1), suggesting that no replicon type is specialized for, or predominately associated with, a particular function. Instead, core cellular functions appear to be distributed across replicon types, suggesting a high degree of functional integration within multipartite archaeal genomes. However, functional composition varied with replicon size, with a clear tendency towards a chr1 functional composition as secondary replicon size increases (Fig. 2). Interestingly, this pattern is not only restricted to replicon length —a threshold feature—, but it is also observable in the progressive increase in TNF similarity —an amelioration feature (Fig. S3). These patterns could result from genetic interchange among replicons within the same genome, as larger secondary replicons tend to share more protein families with chr1 and have more partition-related genes (Fig. 3A, Table S1). The distribution of essential functions across replicon types, together with the increasing homogenization of COG functional categories profiles as replicon length increases, may enhance the long-term functional robustness of multipartite archaeal genomes, and promote the maintenance of a stable genomic network over evolutionary time.

Consistent with the functional annotation results, the pangenome analysis performed at the family level revealed that core, soft-core, accessory, rare, and singleton protein families are distributed across all three replicon types across all examined genomes (Fig. 4A). However, small replicons were enriched in singleton protein families, consistent with rapid gene turnover and the acquisition of lineage-specific functions (Brockhurst et al., 2019), whereas larger replicons, particularly chr1, contained proportionally more core and soft-core protein families (Fig. 4B). Interestingly, accessory protein families reached their highest relative abundance in intermediate-sized replicons (Fig. 4B), suggesting that newly acquired genes may initially persist as accessory components within the pangenome before either becoming fixed through selection and transitioning into the core genome or being lost over evolutionary time (Iranzo et al., 2019).

Collectively, our findings suggest a gradual increase in the functional integration of secondary replicons into the host genome. As genes are selectively retained and their persistence across the replicons in each genome increases, secondary replicons appear to transition from being enriched in rare and accessory functions to harbouring more stable repertoires dominated by core genes. This pattern is consistent with current evolutionary frameworks of pangenome evolution, in which newly acquired genes are continuously gained and tested by selection, eventually becoming broadly conserved across lineages or being lost over evolutionary time (McInerney et al., 2017; Cummins et al., 2022). The progressive stabilization of GC content, TNF, together with the increasing chromosome-like functional composition of larger replicons, supports the existence of an evolutionary continuum between secondary replicons and primary chromosomes.

### Shared functions between chromosomes and plasmids in Halobacteriota are genome-specific

The analysis of the nine multipartite genomes with an annotated chr2 further highlighted the important role of secondary replicons in multipartite archaeal genomes. The extensive sharing of protein families among replicons within the same genome indicates that multipartite archaeal genomes function as integrated genomic systems rather than collections of independent replicons. The central position of chr1 and the high connectivity of chr2 suggest that chr2 are functionally incorporated into the genomic network, whereas plasmids exhibit more variable degrees of integration across species (Fig. S5). However, we observed exceptions. In *Halorubrum lacusprofundi* and *Haloarcula*, chr1 shared more protein family clusters with a replicon annotated as plasmid (n = 30 and n = 72, respectively) than with chr2 (n = 19 and n = 22, respectively) (Fig. S5, Table S7). These cases suggest that chr2 does not always play a role as an intermediate replicon between chr1 and plasmids, as has been proposed for bacteria (Ostermayer et al., 2025), highlighting the diversity of multipartite genome organization in Archaea. However, these cases could also reflect potential misclassification of secondary replicons as chr2 or plasmids.

The greatest differences in GC content among shared protein families located on different replicons were observed in protein families shared only among plasmids, consistent with the greater propensity of plasmids for horizontal gene transfer compared with chromosomes (Sheppard et al., 2020; Rodríguez-Beltrán et al., 2021). The relatively small differences in GC content observed in this analysis provide little support for a recent HGT origin of these shared proteins (Ravenhall et al., 2015). Instead, the HGT events may have occurred sufficiently long ago for compositional differences to have been eroded through amelioration (Lawrence and Ochman, 1997; Lawrence, 2002), or they may have occurred recently from donor replicons with GC content similar to that of the host. The latter explanation is plausible given the hypersaline environments in which Halobacteriota thrive, which may restrict the diversity of microorganisms inhabiting these environments and consequently, limit gene exchange with microorganisms with contrasting GC content. Besides HGT, gene duplication could explain shared protein families with similar GC content. The proportion of duplicated protein families increased with replicon size, supporting gene duplication as a potential mechanism underlying protein-family sharing across replicons (Fig. 3B).

The overall distribution of functional categories was broadly similar across replicon types, however, the functions represented among the shared protein families were not random. Instead, different functional classes exhibit distinct patterns of sharing between replicons (Table S7). Housekeeping functions are preferentially shared between chromosome-like replicons, whereas regulatory proteins and mobile genetic elements are more frequently shared between chr1-plasmid and plasmid-plasmid. Transport systems, in contrast, are shared across all replicon types, suggesting that some functional classes maintain stronger connectivity across genomic compartments than others.

Overall, our analyses suggest that multipartite archaeal genomes do not consist of independent genetic compartments with fixed functional specializations. Instead, they appear to operate as integrated genomic systems in which essential cellular processes are distributed across multiple replicons suggesting that selection acts at the genome level to maintain a genome-wide functional integration. As a consequence, shared clusters appear to be both function-dependent and lineage-specific, indicating that distinct evolutionary processes shape the retention of different functions across multipartite prokaryotic genomes (Koonin and Wolf, 2008; diCenzo and Finan, 2017; Ostermayer et al., 2025).

### Insights into the evolutionary history of archaeal secondary replicons

The similarity in GC content, TNF, and codon usage observed between secondary replicons and chr1 further suggest that secondary replicons are long-term components of the genome rather than recent acquisitions, having undergone extensive compositional amelioration and prolonged co-evolution with their host genomes. Their evolutionary origins are likely diverse, ranging from plasmid acquisition followed by amelioration and domestication (diCenzo and Finan, 2017), to gene gain, or even possible chromosome fission events generating more than one stable replicon (Ausiannikava et al., 2018).

Functional composition further supports the existence of a continuum in replicon organization in Halobacteriota. Across the nine selected genomes analyzed in detail, chr1–chr2 shared families were generally enriched in conserved cellular functions, whereas plasmid-associated families were more often associated with adaptive and mobile functions. Across the full Halobacteriota dataset, replicon size was strongly associated with functional composition: larger secondary replicons contained greater proportions of conserved protein families and showed increased overlap with chr1, whereas smaller replicons were dominated by rare and lineage-specific families. Importantly, pangenome categories were not restricted to particular replicon types, indicating that functional differentiation is gradual rather than discrete. Together, these results suggest that secondary replicons progressively acquire more conserved, chromosome-like gene repertoires as their size increases, while still retaining lineage-specific functional signatures.

Replicon size was also positively associated with protein-family duplication, consistent with previous studies linking genome expansion to increased gene duplication and gene-family expansion (Hooper and Berg, 2003; Manzano-Morales and Gabaldón, 2026). This relationship suggests that duplication may contribute to the expansion of Halobacteriota replicons, while the pangenome analysis provides an additional perspective on the fate of this expanded gene content. Specifically, increasing replicon size was associated with a progressive decline in singleton and rare protein families and an increase in soft-core and core families (Fig. 4B). This pattern is consistent with differential retention of genes, in which some genes are maintained across lineages while others are lost. Thus, replicon expansion appears to involve not only the acquisition and duplication of genes, but also their subsequent retention and integration over evolutionary time.

Although the precise evolutionary trajectories likely differ among archaeal taxonomic families, the combined evidence presented here supports a scenario in which ancient replicon acquisition, differential retention, gene duplication, and long-term selection progressively increase the structural, compositional and functional integration of secondary replicons into the host genome. Rather than persisting as largely independent genetic elements, successful secondary replicons appear to become stable components of multipartite archaeal genomes through the gradual stabilization of their gene repertoires and their increasing contribution to genome-wide cellular functions.

These patterns should nevertheless be interpreted with caution given the limited availability of complete multipartite Halobacteriota genomes. Although 211 genomes were included in this study, secondary replicons recognized as chr2 replicons were found in only nine genomes, limiting the taxonomic breadth available for comparisons involving secondary chromosomes. As additional complete genomes become available, particularly from currently underrepresented lineages, it will be possible to determine how consistently the genomic and functional patterns described here extend across Halobacteriota and in Archaea more broadly.

## CONCLUSIONS

Taken together, our findings suggest that secondary replicons occupy different stages along an evolutionary continuum of genomic integration, rather than representing discrete genomic entities. Replicons at early stages of this continuum are expected to exhibit greater compositional divergence from chr1, together with an enrichment of accessory and lineage-specific genes. Over evolutionary time, compositional amelioration, the differential retention of beneficial functions, and the expansion of gene families through duplication may progressively increase the structural, compositional and functional integration of secondary replicons into the host genome. Consistent with this observation, larger secondary replicons contained a higher number of duplicated protein families, exhibited a progressive shift from singleton and accessory toward persistent and core gene families, shared a greater proportion of housekeeping functions with chromosome-like replicons, and generally showed lower compositional divergence. Remarkably, these independent lines of evidence, including sequence composition, pangenome dynamics, gene duplication, and functional organization, converge on the same evolutionary interpretation, supporting the existence of a continuum of secondary replicon integration in the Halobacteriota. Within this model, secondary replicons —both chr2 and plasmids— may represent highly integrated secondary replicons that have acquired essential cellular functions while retaining signatures of their distinct evolutionary origins, effectively bridging the continuum between mobile plasmids and chr1.

## ACKNOWLEDGEMENTS

This work was funded by the Consejo Superior de Investigaciones Científicas (CSIC), grant (20243MAX009) (to M.T.-R).

