## Supplementary Material for "Comparative genomics reveals a genomic and functional continuum among secondary replicons in Halobacteriota"

### Supplementary figures

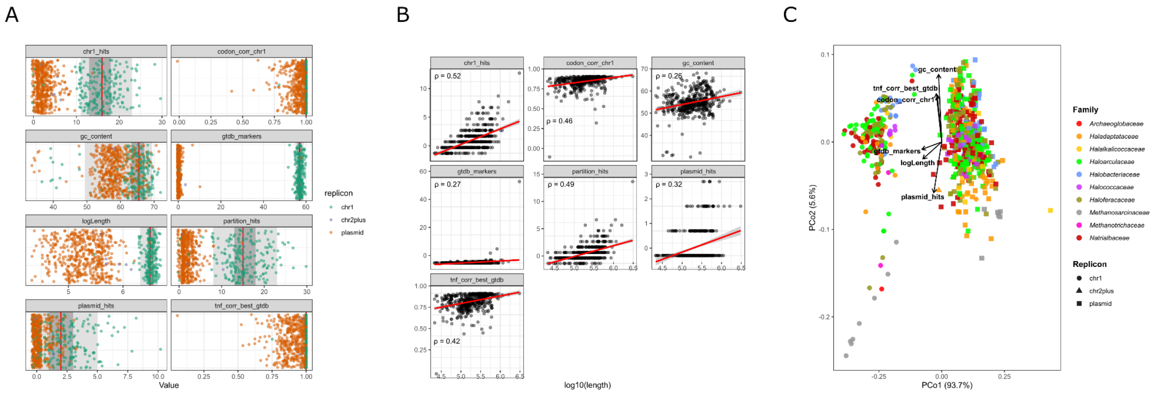


**Figure S1. Genomic landscape and feature changes according to length.** (A) Distribution of all replicons (including TNF and Codon Usage outliers) in the dataset according to their value in each of the evaluated features. Each x-axis corresponds to the range of values associated with each of the features. (B) Dispersion plots showing the values of each feature according to length. Each point represents a replicon and Spearman's rank correlation coefficient (ρ) is shown. (C) PCoA ordination showing all replicons in the Halobacteriota dataset. Each point corresponds to a replicon, color indicates taxonomic family, and shape indicates replicon type. Vectors corresponding to the different genomic features are shown. Statistical significance was evaluated through PERMANOVA, $R^2 = 0.853$, $F = 2117.51$, $P < 0.0001$, and PERMDISP, $F = 6.58$, $P = 0.006$.

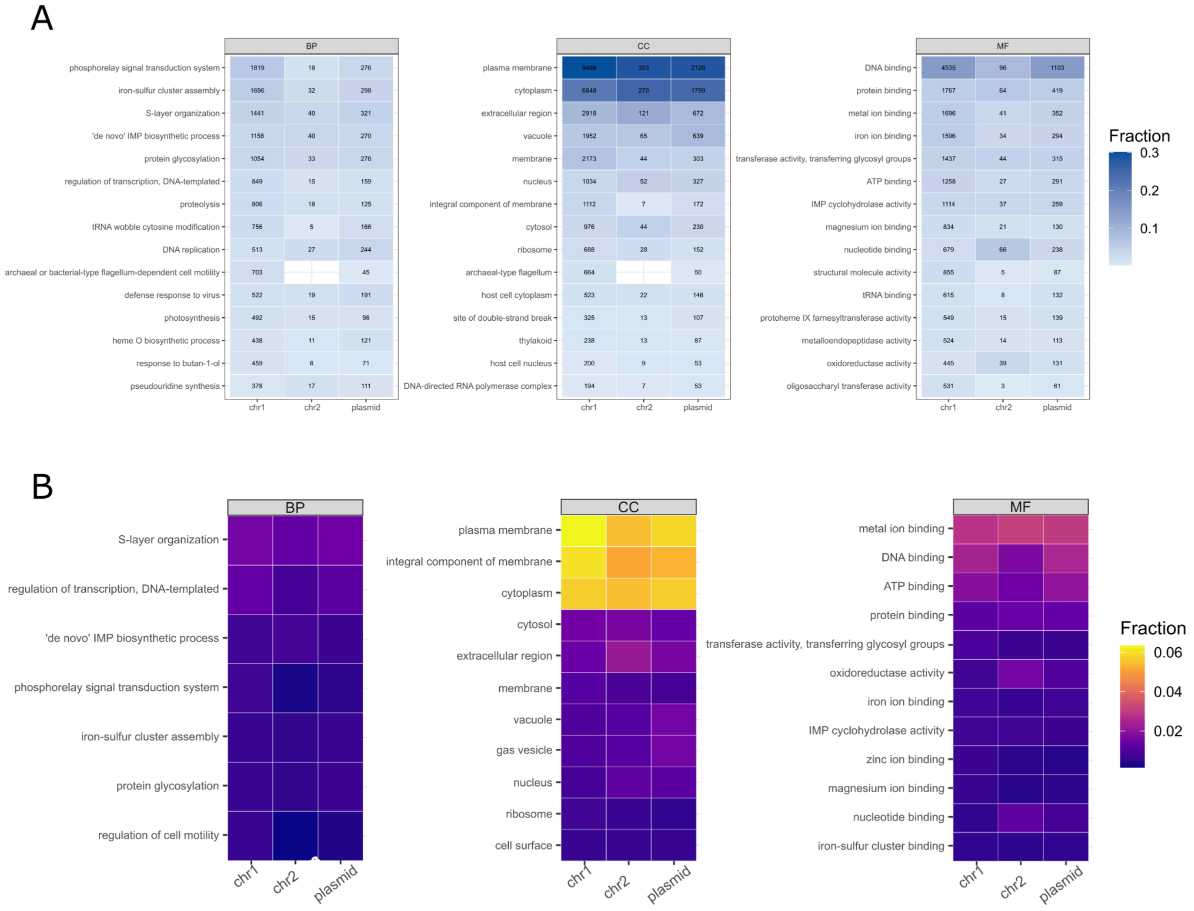


**Figure S2. Distribution per replicon type of unknown protein families.** (A) Distribution of the top 15 recovered functions across replicons, shown as the fraction of clusters assigned to each function within each ontology and replicon. Numbers indicate the number of clusters assigned to each function. (B) Distribution of Gene Ontology (GO) terms among unknown proteins across replicons, showing the fraction of GO-annotated proteins associated with each of the top 30 GO terms within each replicon. GO terms are grouped by ontology.

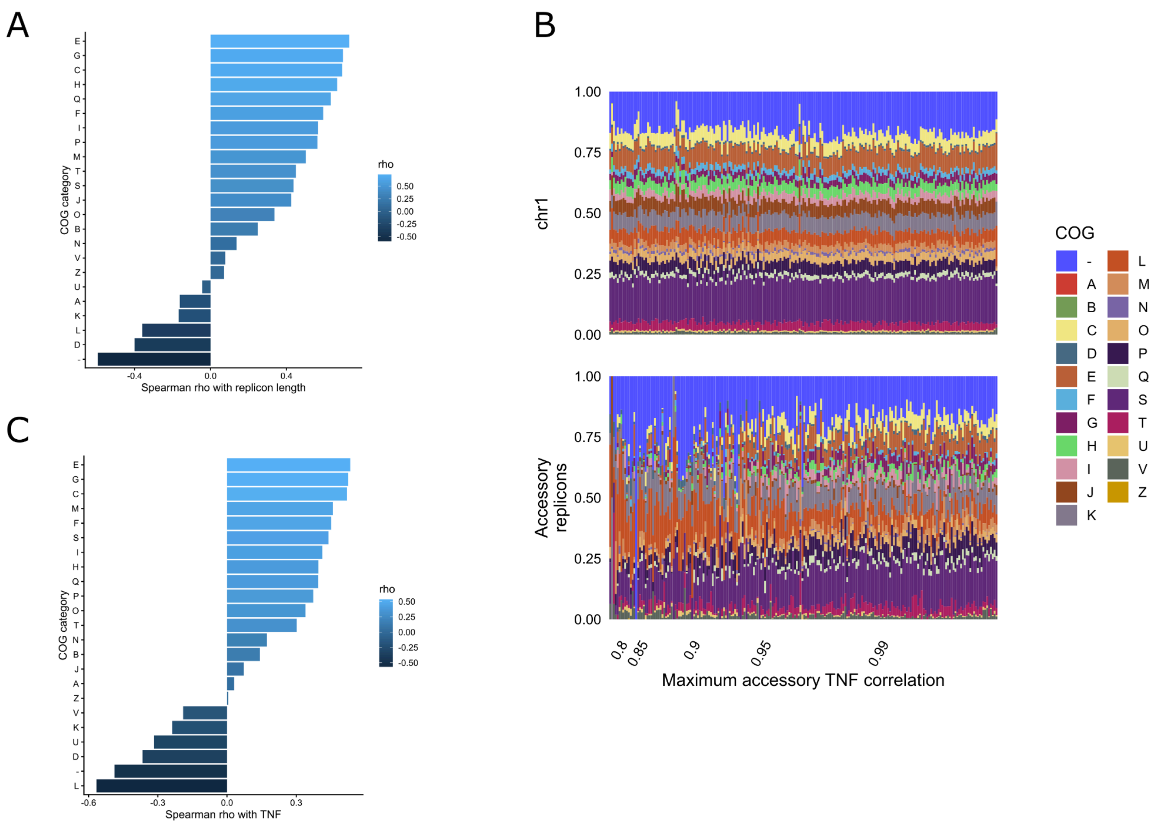


**Figure S3. Chromosome-like features increase with TNF value.** (A) Bar plot of COG categories correlated with length according to Spearman ρ value. Categories with values under zero are negatively correlated, while categories with values above zero are positively correlated. (B) Barplots showing COG functional content per replicon. The lower plot shows secondary replicons ordered by increasing length, with the corresponding chr1 replicons shown in the upper barplot according to the ordering of their associated secondary replicons. COG functional content of the lower plot becomes remarkably similar to the pattern in the upper plot as replicon length increases. (C) Dispersion plots showing the relative abundance of each COG category per replicon according to TNF values. Each point represents a replicon and fitted curves are shown.

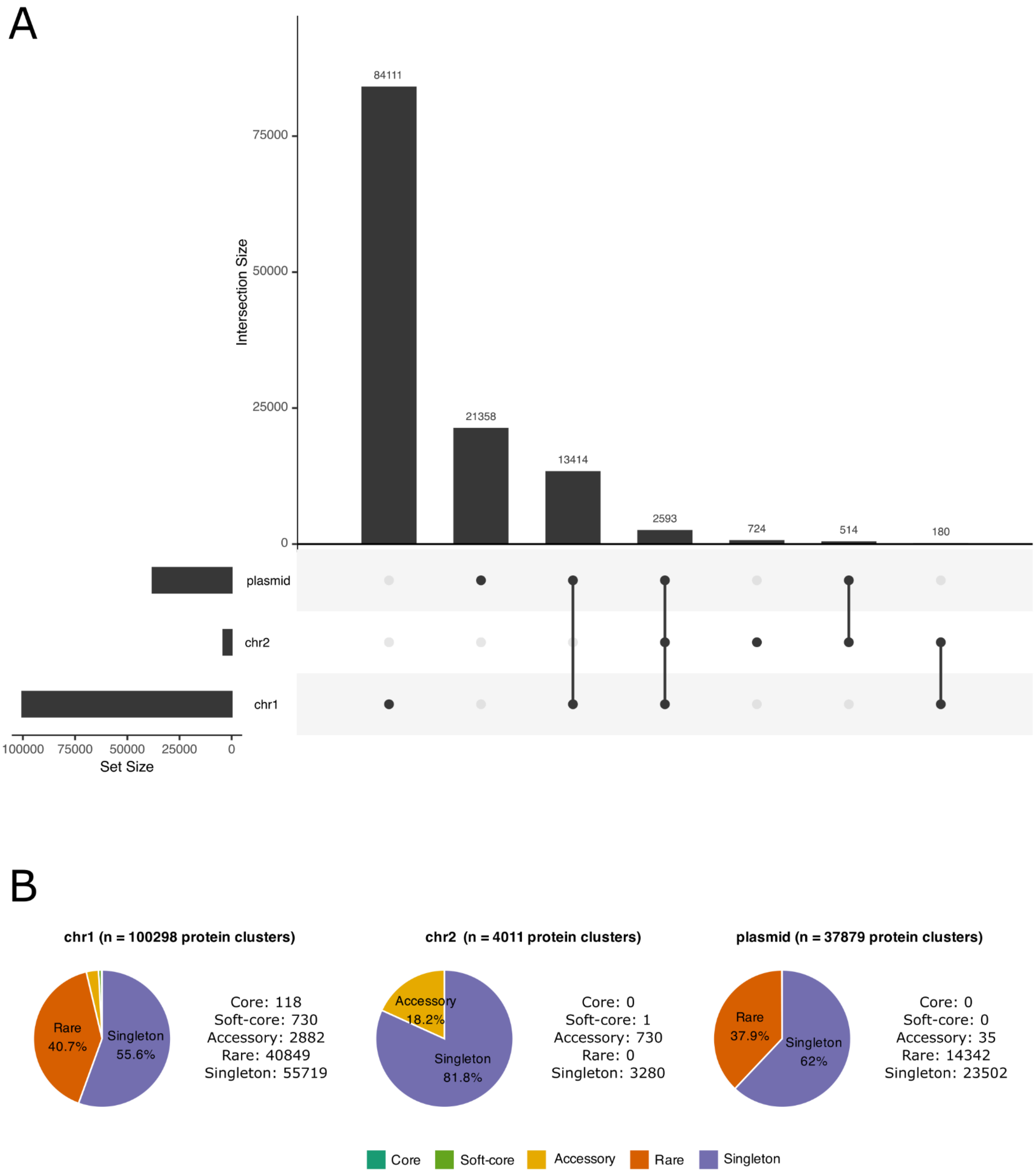


**Figure S4. Halobacteriota pangenomes.** (A) UpSetR plot depicting the number of protein families shared among the different replicon types. This analysis was conducted using replicon type as comparison units and ignoring their taxonomical origin. (B) Pie charts representing the number (raw counts and percentages) of proteins per pangenome category (core, soft-core, accessory, rare, and singletons). Each pie chart represents the pangenome constructed using only chr1 (left), chr2 (middle), and plasmids (right).

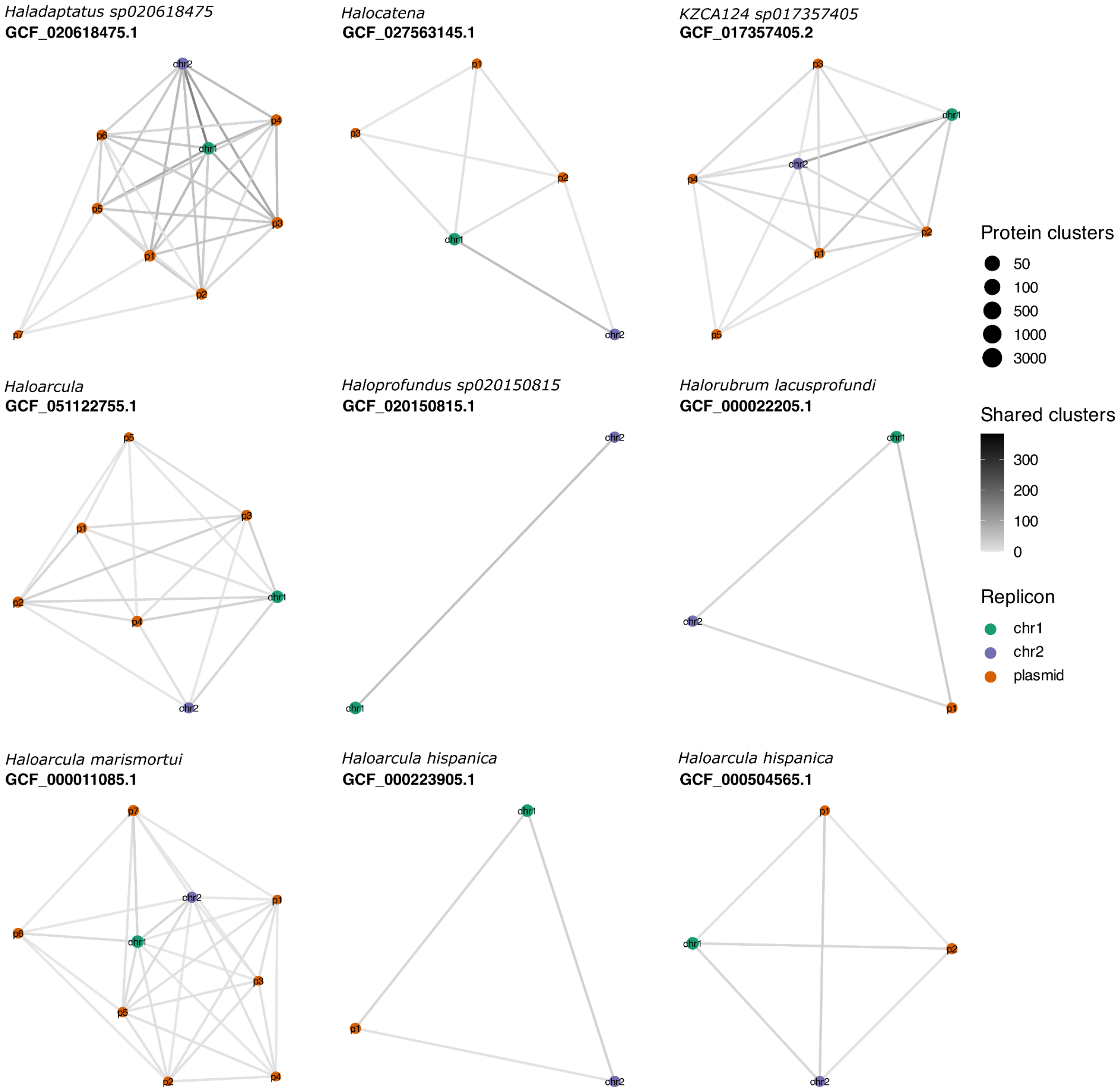


**Figure S5. Shared protein families between the replicons present in nine different genomes.** A network is shown per each of the nine genomes analyzed. Nodes represent replicons colored according to replicon type, with size representing the number of clusters contained in each replicon. Edges represent shared protein families between the connected nodes, with a grey gradient indicating the number of shared clusters. The corresponding taxonomic annotation along with the accession number are indicated above each one of the networks.

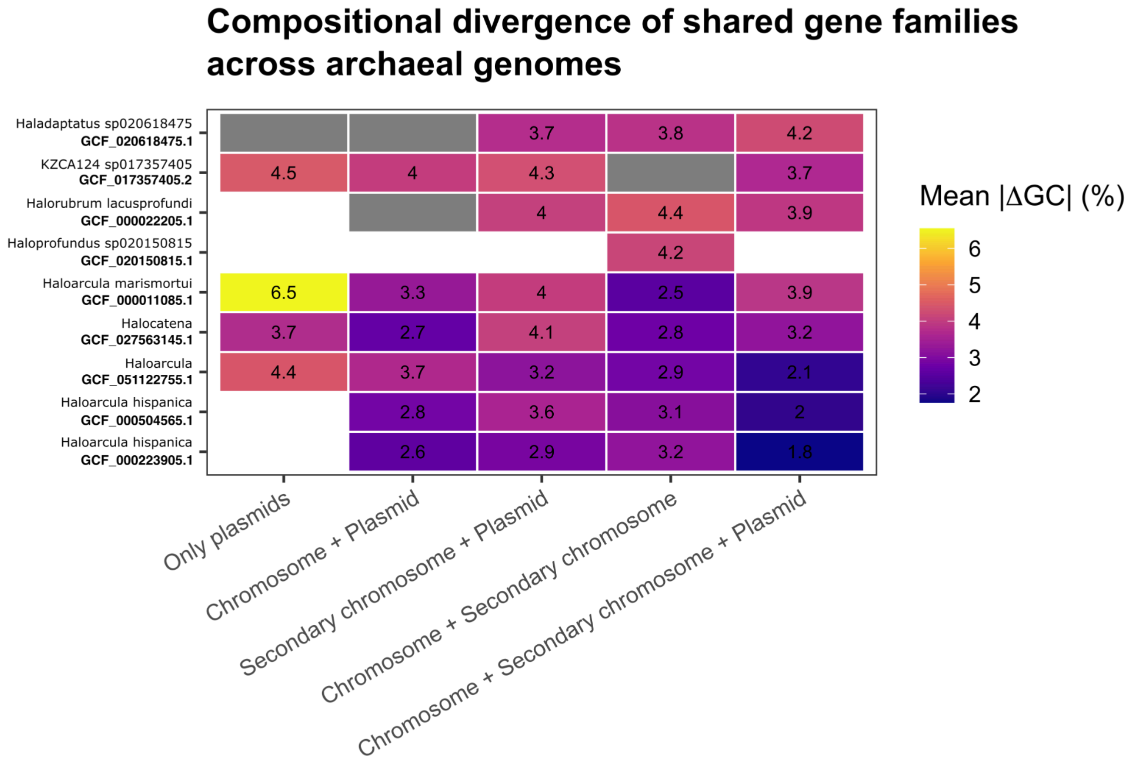


**Figure S6. Differences in GC content of protein families shared among different replicon types per genome.** Mean |ΔGC| for each genome and shared-cluster category obtained from the summatory of the absolute GC-content difference (|ΔGC|). Higher values in mean |ΔGC| suggest larger GC content differences among the shared protein families among the different replicon pairs.

### Supplementary tables

All supplementary tables are available on GitHub
https://github.com/ajaraservin/archaea_secondary_replicons

**Table S1.** Genomic features and taxonomic annotation of all the genomes used in this study.

**Table S2.** Families and their features in upper and lower clusters.

**Table S3.** Functional annotation of archaeal protein families using COG database

**Table S4.** FANTASIA functional annotation of S COG category and COG non-annotated archaeal protein families.

**Table S5.** Functional annotation of shared and unique protein families according to replicon type from the UpSetR analysis.

**Table S6.** Protein families comprising the chr2 pangenome.

**Table S7.** Functional annotation of all shared protein families per replicon per genomes harboring a chr2.

**Table S8.** Annotation of shared protein families between replicons per genome using FANTASIA.
